# A community-based collaboration to build prediction models for short-term discontinuation of docetaxel in metastatic castration-resistant prostate cancer

**DOI:** 10.1101/087809

**Authors:** Fatemeh Seyednasrollah, Devin C Koestler, Tao Wang, Stephen R Piccolo, Roberto Vega, Russ Greiner, Christiane Fuchs, Eyal Gofer, Luke Kumar, Russell D Wolfinger, Kimberly Kanigel Winner, Chris Bare, Elias Chaibub Neto, Thomas Yu, Liji Shen, Kald Abdallah, Thea Norman, Gustavo Stolovitzky, PCC-DREAM Community, Howard Soule, Christopher J Sweeney, Charles J Ryan, Howard I Scher, Oliver Sartor, Laura L Elo, Fang Liz Zhou, Justin Guinney, James C Costello

## Abstract

**Background:** Docetaxel has a demonstrated survival benefit for metastatic castration-resistant prostate cancer (mCRPC). However, 10-20% of patients discontinue docetaxel prematurely because of toxicity-induced adverse events, and managing risk factors for toxicity remains an ongoing challenge for health care providers and patients. Prospective identification of high-risk patients for early discontinuation has the potential to assist clinical decision-making and can improve the design of more efficient clinical trials. In partnership with Project Data Sphere (PDS), a non-profit initiative facilitating clinical trial data-sharing, we designed an open-data, crowdsourced DREAM (Dialogue for Reverse Engineering Assessments and Methods) Challenge for developing models to predict early discontinuation of docetaxel

**Methods:** Data from the comparator arms of four phase III clinical trials in first-line mCRPC were obtained from PDS, including 476 patients treated with docetaxel and prednisone from the ASCENT2 trial, 598 patients treated with docetaxel, prednisone/prednisolone, and placebo in the VENICE trial, 526 patients treated with docetaxel, prednisone, and placebo in the MAINSAIL trial, and 528 patients treated with docetaxel and placebo in the ENTHUSE 33 trial. Early discontinuation was defined as treatment stoppage within three months due to adverse treatment effects. Over 150 clinical features including laboratory values, medical history, lesion measures, prior treatment, and demographic variables were curated and made freely available for model building for all four trials. The ASCENT2, VENICE, and MAINSAIL trial data sets formed the training set that also included patient discontinuation status. The ENTHUSE 33 trial, with patient discontinuation status hidden, was used as an independent validation set to evaluate model performance. Prediction performance was assessed using area under the precision-recall curve (AUPRC) and the Bayes factor was used to compare the performance between prediction models.

**Results:** The frequency of early discontinuation was similar between training (ASCENT2, VENICE, and MAINSAIL) and validation (ENTHUSE 33) sets, 12.3% versus 10.4% of docetaxel-treated patients, respectively. In total, 34 independent teams submitted predictions from 61 different models. AUPRC ranged from 0.088 to 0.178 across submissions with a random model performance of 0.104. Seven models with comparable AUPRC scores (Bayes factor ≤; 3) were observed to outperform all other models. A post-challenge analysis of risk predictions generated by these seven models revealed three distinct patient subgroups: patients consistently predicted to be at high-risk or low-risk for early discontinuation and those with discordant risk predictions. Early discontinuation events were two-times higher in the high-versus low-risk subgroup and baseline clinical features such as presence/absence of metastatic liver lesions, and prior treatment with analgesics and ACE inhibitors exhibited statistically significant differences between the high- and low-risk subgroups (adjusted *P* < 0.05). An ensemble-based model constructed from a post-Challenge community collaboration resulted in the best overall prediction performance (AUPRC = 0.230) and represented a marked improvement over any individual Challenge submission. A

**Findings:** Our results demonstrate that routinely collected clinical features can be used to prospectively inform clinicians of mCRPC patients’ risk to discontinue docetaxel treatment early due to adverse events and to the best of our knowledge is the first to establish performance benchmarks in this area. This work also underscores the “wisdom of crowds” approach by demonstrating that improved prediction of patient outcomes is obtainable by combining methods across an extended community. These findings were made possible because data from separate trials were made publicly available and centrally compiled through PDS.

## Introduction

The long-term prognosis of metastatic castration-resistant prostate cancer (mCRPC) is poor with median overall survival ranging on average, from 10 to 27 months, depending on metastatic site(s)^1^. Docetaxel was the first cytotoxic drug to improve mCRPC survival and quality of life^2, 3^, and has remained the standard first-line chemotherapy for treating mCRPC. Although several clinical trials have since confirmed docetaxel’s population-level survival and palliative benefits^4, 5^, a significant fraction of patients do not respond to docetaxel and within approximately 8 months, nearly all patients become resistant or have stopped therapy due to toxicity^2, 3^. Additionally, of those initially responding to docetaxel, 10-20% prematurely discontinue due to toxicity-induced adverse events (AE) that include anemia, (febrile) neutropenia, fatigue, fluid retention, nail toxicity, gastrointestinal complications, and neuropathies^6–8^. Managing risk factors for toxicity is a major challenge for health care providers as they may hinder patients from receiving a therapy with potential clinical benefit, and/or diminish a patient’s quality of life without extending life.

As docetaxel-based chemotherapy continues to play an important role in the treatment of mCRPC and more recently hormone sensitive metastatic prostate cancer^9^, it is important to prospectively identify patients for whom a docetaxel-based regimen is likely to be poorly tolerated, resulting in AE and potentially early treatment failure. In particular, such knowledge could be used to pinpoint patients for preemptive clinical interventions/supportive care prior to chemotherapy, when such measures are likely to be most effective, or direct patients to alternative treatment regimens. In addition, establishing quantitative benchmarks for identifying patients at high-risk for early docetaxel discontinuation can be used to facilitate the design of more efficient trials by assisting the selection of a more homogenous patient populations. Finally, identifying and precluding patients who are likely to have adverse response(s) provides an ethical advantage over clinical trials that make no such distinction. Although prognostic models in mCRPC have been previously described^10–13^, there are currently no companion quantitative tools that facilitate prospective risk predictions for early treatment discontinuation based on a patient’s unique clinical characteristics.

Here, we report the results from the Prostate Cancer DREAM (Dialogue for Reverse Engineering Assessments and Methods) Challenge, the first crowdsourced competition in mCRPC with the aim to improve predictions of toxicity in docetaxel-treated mCRPC patients. This Challenge builds on the open clinical trial data initiative of Project Data Sphere LLC (PDS) - a non-profit initiative of the CEO Roundtable on Cancer’s Life Consortium. The comparator arms of four, phase III clinical trials were made public, representing a major contribution that removed the privacy and legal barriers for open-data access. Three of the four trials formed the training data set (*n* = 1,600) and the fourth trial data set (*n* = 470) was used for independent evaluation and validation of model prediction performance. Over 150 clinical features were made available for the trials. Over a five-month competition period, 34 teams from around the world worked independently to address the challenge of predicting early discontinuation of docetaxel due to AE. We present novel clinical variables that are associated with treatment discontinuation and provide a statistical analysis of clinical trial designs that incorporate likelihood of discontinuation in the patient selection criteria. Finally, we describe a post-Challenge community-based collaboration between Challenge organizers and participating teams members – individuals that had never before collaborated prior to this Challenge – aimed at leveraging the “wisdom of crowds” to further refine risk-prediction models.

## Methods

### Trial selection, patient population, and data processing

In April 2014, the data used in this challenge were complied based on de-identified comparator arm data sets of four Phase III prostate cancer clinical trials hosted on *Project Data Sphere* (PDS). All four trials (ASCENT2^14^, VENICE^15^, MAILSAIL^16^, and ENTHUSE 33^17^) were randomized and shared similar inclusion/exclusion criteria; eligible patients included those with progressive mCRPC, no previous chemotherapy, and an Eastern Cooperative Oncology Group (ECOG) performance status of 0 to 2. Detailed inclusion/exclusion criteria of each trial can be found in the Supplementary Appendix. These patient-level trial datasets were de-identified by data providers and made available for the Challenge through PDS. In total, the data used in this Challenge consisted of 2,070 first-line mCRPC patients treated with a docetaxel-based treatment regimen, enrolled in one of the following trials:

#### ASCENT2^14^ (Novacea, provided by Memorial Sloan Kettering Cancer Center)

ASCENT2 is a randomized, open-Label study evaluating DN-101 in combination with docetaxel in mCRPC. Patients received docetaxel and prednisone in the comparator arm (*n* = 476; 105 patients discontinued docetaxel within three months, due to AE or possible AE).

#### VENICE^15^ (Sanofi)

VENICE is a randomized, double-blind study comparing efficacy and safety of aflibercept versus placebo in mCRPC patients treated with docetaxel and prednisone. Patients received docetaxel, prednisone and placebo in the comparator arm (*n* = 598; 51 patients discontinued docetaxel within three months, due to AE or possible AE).

#### MAINSAIL^16^ (Celgene)

MAINSAIL is a randomized, double-blind study to evaluate efficacy and safety of docetaxel and prednisone with or without lenalidomide among mCRPC patients. Subjects received docetaxel, prednisone and placebo in the comparator arm (*n* = 526; 41 patients discontinued docetaxel within three months, due to AE or possible AE).

#### ENTHUSE 33^17^ (AstraZeneca)

ENTHUSE 33 is a randomized, double-blind study to assess efficacy and safety of 10 mg ZD4054 combined with docetaxel in comparison with docetaxel only among mCRPC patients. Subjects received docetaxel and placebo in the comparator arm (*n* = 470; 49 patients discontinued docetaxel within three months, due to AE or possible AE).

The ASCENT2, VENICE, and MAINSAIL data sets were combined to create the training data set (*n* = 1,600) and the ENTHUSE 33 data defined the independent validation set to evaluate model prediction performance. Due to regulation and privacy restrictions of certain countries, data from 470 patients in the comparator arm of ENTHUSE 33 (*n* = 528 in total) were provided to PDS. Additional details describing data splitting into training and validation sets is given in the Supplementary Appendix.

### Data curation

The original data sets from PDS contained patient level raw tables that conformed to either Study Data Tabulation Model (SDTM) standards or company-specific clinical database standards. The four sets of raw trial data were consolidated into set of five standardized raw even-level tables covering lab values, medical history, lesion measures, prior therapies, and vital signs. The standardized raw even-level tables, including patient demographics comprised more than 150 variables of potential clinical importance. The raw event-level tables were then summarized for each individual patient into a “Core Table” representing a total of 129 baseline and outcome variables. The five raw event-level tables and the Core Table were made available to teams for each of the trials. Patient discontinuation status was withheld from the validation data set (ENTHUSE 33 trial). Full details of data curation can be found in the Supplementary Appendix.

### Creation of the dependent variable

The dependent variable (DISCONT) was derived from two factors: reason for treatment discontinuation (i.e., “discontinue reason”) and the time from treatment initiation to discontinuation (i.e., “discontinue time”). Discontinuation of treatment was evaluated for the first 3 months of treatment, or the first 4 cycles (12 weeks) of treatment in a 10 cycle regimen, 3 weeks per cycle. Reasons for treatment discontinuation were grouped into five major categories: 1) discontinuation due to an AE, 2) discontinuation possibly due to an AE, 3) death or progression, 4) completed treatment, and 5) a miscellaneous group (Table S1 and Supplementary Appendix). Patients were labeled as DISCONT=1 if and only if they discontinued treatment due to AE or possible AE within 3 months (91.5 days) after beginning treatment, otherwise patients were labeled as DISCONT=0. The number and percentage of patients assigned to each the above categories are given in Table S2.

### Challenge design, scoring, and evaluation

The Challenge was hosted on Synapse (<u>www.synapse.org</u>), a free, cloud-based platform for collaborative scientific data analysis. Synapse was used to allow access to Challenge data and to track participant agreements to the appropriate data use agreements (<u>https://www.synapse.org/#!Synapse:syn3348040</u>) and Challenge rules (<u>https://www.synapse.org/#!Synapse:syn3348041</u>).

Teams were tasked with developing models to predict early discontinuation of docetaxel due to AE or possible AE. Six-weeks prior to the Challenge deadline, teams were given access to the patient-level clinical data for the validation set (Fig. S1). Using these data, teams submitted up to two risk scores (i.e. predictions) for each patient. For final submissions, Challenge participants were required to create open-access Synapse projects containing their predictions, corresponding code, and a write-up describing their analytical approach. Risk scores submitted by each team were subsequently evaluated and ranked using the area under the precision-recall curve (AUPRC)^18^. AUPRC values range between 0 and 1, with larger values indicating better prediction performance. The AUPRC was selected over the more commonly used area under the receiver operating characteristic curve (AUROC) because the dataset is imbalanced with 10-20% of patients discontinuing treatment. We focus on a method’s ability to predict the discontinued patients (positive cases) and not the overall performance; the AUROC does not properly calculate the performance of predicting positive cases in unbalanced data. For teams that submitted two prediction models, the larger of the two AUPRCs was used to determine their final placement in the leaderboard. Since 10.4% of patients in ENTHUSE 33 were labeled as DISCONT = 1, the expected AUPRC for a random prediction model is 0.104; only submissions that exceeded this threshold were considered to provide potential clinical value.

The following criteria were used to determine the top teams/models: (1) prediction performance was significantly better than a random prediction model and (2) performance was statistically indistinguishable when compared to the model achieving the highest AUPRC score. To assess whether a model’s prediction performance was significantly better than random, its AUPRC was compared to the empirical null distribution, generated from 5,000 random permutations of the dependent variable^19^. One-sided p-values were computed as the probability of observing an AUPRC under the null distribution that was at least as large as the AUPRC obtained for a given team. P-values were corrected for multiple testing using the Benjamini-Hochberg procedure^20^, and adjusted p-values less than 10% (*P* < 0.10) were considered statistically significant. To assess whether consecutively ranked models were measurably distinguishable in terms of their AUPRC score, the Bayes factor^21, 22^ was computed between each model and the first ranked model. Submissions with a Bayes factor ≤ 3 from the first ranked model were declared statistically indistinguishable. Further details concerning model scoring and evaluation can be found in the Supplementary Appendix.

As an alternative to AUPRC and to provide additional insight into the clinical utility of prediction models, risk scores submitted by each team were subjected to a cumulative lift chart analysis (Supplementary Appendix). For each team, results were summarized by computing: (1) the area under the lift ratio curve and (2) the lift ratio evaluated among patients with the highest predicted for early treatment discontinuation risk (top 5%, 10%, and 20%).

Principal Component Analysis (PCA) was conducted to explore systematic similarities or differences between the four studies. PCA was conducted either using all available variables or only using binary variables. Visualization of PCA was done by plotting the first principal component against the second principal component for all patients.

Hierarchical clustering was performed using Ward’s method and Manhattan distance.

### Post-Challenge community collaboration to improve patient risk predictions

Following the completion of the Challenge, an ensemble-based prediction model^23^ was generated using the top seven teams’ models (Fig. S2 and Supplementary Appendix). To construct the ensemble-based model, top performing teams ran their model *L*_*i*_(⋅), *i* = 1, … *P*, on the full training data *D* to produce the following predictors: {*C*_1_(*r*),…, *C*_P_(*r*)}; *P* denotes the number of top teams/models identified from the Challenge and *C*_*i*_(*r*) represents the estimated risk of early treatment discontinuation for patient *r* based on the *i*^th^ model. Using the predictors generated by each of the top teams/models, an ensemble-based prediction model was generated as the following simple, weighted average:

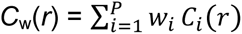

with weights, *w*_*i*_ *i* = 1, …, *P*, proportional to the prediction accuracy of *C*_*i*_(.). To learn these weights, the training data, *D*, was randomly split into two independent sets: *D*^*70*^, which contained 70% of the patients in the training data (*n* = 1,120) and *D*^*30*^, which contained the remaining 30% (*n* = 480). Models, *L*_*i*_(⋅), developed by the top teams were first trained on *D*^*70*^ to produce seven new predictors: {*C*_*i*_^*70*^(.), *i* = 1, …, *P*}. Each of these predictors were then used to predict the early discontinuation status (i.e., = 1 early discontinuation; = 0 otherwise) for each patient in *D*^*30*^. Because the outcome of interest (i.e., early discontinuation status) was observed for all patients in *D,* and consequently *D*^*30*^, the prediction accuracy associated with each of the classifiers, *A*_*i*_, was computed as the fraction of patients that were correctly predicted to prematurely discontinue treatment. With weights set to *w*_*i*_ = *A*_*i*_, the ensemble-based prediction model, *C*_*w*_(*r*), was applied to the ENTHUSE 33 data and its AUPRC was computed. To determine if the AUPRC score generated from the ensemble-based prediction model represented an improvement over the scores obtained from individual model submissions, bootstrap sampling was used to approximate the distribution of AUPRC for each team, as well as for the ensemble-based model. For each bootstrap sample (5000 total replications), the difference in the AUPRC scores between the ensemble-based model and individual submissions were computed, allowing us to estimate the fraction of times the ensemble-based model outperformed each of the individual model submissions. The Bayes factor between each team and the ensemble-based model was also calculated using the procedure described above.

As there was no restriction imposed by Challenge organizers on the number of clinical features used in the development of prediction models, several of the prediction models submitted to the Challenge - including the post-Challenge ensemble-based model described above - used many or all of the baseline clinical features contained in the standardized data table. In an effort to develop a more parsimonious prediction model (i.e., one using a limited number of baseline clinical features), we developed a second prediction model using only those clinical features that best discriminated subjects predicted to have a high-versus low-risk of early treatment discontinuation using the challenge results; we hereafter refer to this model as the community-based parsimonious prediction model. Briefly, using the risk predictions submitted by each of the top-performing teams for the patients in *D*^*30*^, we performed a hierarchical clustering analysis (Manhattan distance and Ward linkage) to identify patients that were consistently predicted to have a high-risk of early treatment discontinuation versus those predicted to have a low-risk of early treatment discontinuation, across the seven top-performing teams. We next identified baseline clinical features that were significantly different between the high- and low-risk groups by independently testing the association between each baseline clinical feature and patient subgroup using the appropriate univariate test; Wilcoxon rank-sum test for continuous features and a Fisher’s exact test for binary and categorical features. Statistically significant clinical features (*P* < 0.05) identified from this analysis were then carried forward and used to generate a prediction model by fitting a Cox proportional hazards model to the entire training data set *D*, using those features as predictors. Similar to the previously described ensemble-based prediction model, this “community-based parsimonious prediction model” was applied to the ENTHUSE 33 validation set to generate a risk prediction for each of the patients in this data set. Risk predictions were used to compute the AUCPR for comparison with the ensemble-based prediction model and Challenge submissions.

### Clinical trial model simulations

A simulation study was used to compare the sample size requirements of clinical trials that incorporated baseline estimates of a patient’s risk for early treatment discontinuation into patient selection schemes. For our simulation study, we assumed a two-arm randomized controlled trial, 1:1 randomization between arms (i.e. treatment versus control), and survival time as the primary endpoint of interest. The goal of our simulation study was to demonstrate that patient selection schemes that make use of a patient’s baseline line risk for early discontinuation by down-weighting the selection probabilities of “at risk” patients, result in smaller trials without compromising statistical power for detecting the desired effect size.

To simulate realistic survival data, we used the ENTHUSE 33 (validation data) to inform suitable simulation parameters. A parametric survival model (assuming an exponential distribution) was first fit to the ENTHUSE 33 data set and used to estimate the parameters governing the time-to-event and censoring distributions, including the hazard ratio (HR) between docetaxel-treated patients that did and did not discontinue treatment early. Using these parameters, the “survsim” package in R was used to jointly simulate survival data for patients in the treatment and control arm assuming a 10.4% rate of early treatment discontinuation among patients in the treated group, consistent with the discontinuation rate observed in the ENTHUSE 33 data set. In total, 100 independent data sets were simulated, each containing 10,000 patients. Within each of 100 simulated data sets, patients were randomly selected with replacement (1:1 treatment versus control groups) and used to estimate that sample size required for detecting a survival difference (i.e., HR) between the treated and control groups at 80% statistical power and assuming a type 1 error rate of 5%. Patients identified as “at risk” for early discontinuation were excluded from randomization for baseline prediction models with 0%, 25%, 50%, 75%, and 100% accuracy at identifying true cases of early discontinuation.

### Data and method availability

The clinical trial data used in the Challenge can be accessed at <u>https://www.projectdatasphere.org/projectdatasphere/html/pcdc</u>. Method write-ups, code, and predictions for all teams are reported in Tables S3,S4. Documentation, including a detailed description of the Challenge design, overall results, scoring scripts, trial data sets, and data dictionary can be found at: <u>https://www.synapse.org/ProstateCancerChallenge</u>.

## Results

The overall Challenge design is illustrated in Figure 1A. Over 150 baseline clinical variables and longitudinal features comprised the complete aggregated data set, and included: demographic variables, lab values, lesion measurements, medical history, previous medical procedures, and concomitant medications. These variables were harmonized across the four trials to create a single standardized data set, which served as the primary data source for model building and development. Although the majority of baseline clinical variables were fairly consistent across the four trials, notable differences in the distribution of binary clinical features – primarily representing lesion sites – were observed across trial data sets (Table 1, Fig. S3); ASCENT2 patients had much lower percent of visceral metastases (1.1% liver and 1.7% lung) compared to patients in the other three trials (10-14% liver, 11-15% lung). The frequency of early discontinuation events was similar between training and validation sets (12.3% versus 10.4% of treated patients, respectively), but varied considerably across individual trials; ASCENT2 trial had the highest proportion of patients that discontinued treatment within three months (22.1%), followed by ENTHUSE 33 (10.4%), VENICE (8.5%), and MAINSAIL (7.8%) trials (Figure 1C).

**Table 1.** Summary of selected baseline clinical characteristics across trials. Variables that show significant difference between training and validation datasets (K-S test or Chisq test p value<0.05) are marked in *.

| Characteristics | Training Set |  |  | Validation set |
| --- | --- | --- | --- | --- |
|  | ASCENT2<br>(n=476) | MAINSAIL<br>(n=526) | VENICE<br>(n=598) | ENTHUSE 33<br>(n=470) |
| **Age** |  |  |  |  |
| 18-64 | 111 (23.3%) | 171 (32.5%) | 219 (36.6%) | 160 (34.0%) |
| 65-74 | 211 (44.3%) | 246 (46.8%) | 254 (42.5%) | 217 (46.2%) |
| >=75 | 154 (32.4%) | 109 (20.7%) | 125 (20.9%) | 93 (19.8%) |
| **ECOG PS\*** |  |  |  |  |
| 0 | 220 (46.2%) | 257 (48.9%) | 280 (46.8%) | 247 (52.6%) |
| 1 | 234 (49.2%) | 247 (47.0%) | 291 (48.7%) | 223 (47.4%) |
| 2 | 22 (4.6%) | 20 (3.8%) | 27 (4.5%) | 0 (0.0%) |
| **Metastasis** |  |  |  |  |
| Liver* | 5 (1.1%) | 58 (11.0%) | 60 (10.0%) | 64 (13.6%) |
| Bone* | 345 (72.5%) | 439 (83.5%) | 529 (88.5%) | 470 (100%) |
| Lungs | 8 (1.7%) | 74 (14.1%) | 88 (14.7%) | 56 (11.9%) |
| Lymph nodes | 163 (34.2%) | 298 (56.7%) | 323 (54.0%) | 208 (44.3%) |
| **Analgesic use** |  |  |  |  |
| No | 338 (71.0%) | 347 (66.0%) | 419 (70.1%) | 339 (72.1%) |
| Yes | 138 (29.0%) | 179 (34.0%) | 179 (29.9%) | 131 (27.9%) |
| **LDH, U/L** |  |  |  |  |
| 1<sup>st</sup> Quantile | 176 | 174 | NA | 181 |
| Median | 202 | 210 | NA | 213 |
| 3rd Quantile | 250 | 267 | NA | 287 |
| Missing | 13 (2.7%) | 1 (0.2%) | 596 (99.7%) | 5 (1.1%) |
| **PSA, ng/mL** |  |  |  |  |
| 1<sup>st</sup> Quantile | 24.2 | 32.2 | 30.8 | 33.6 |
| Median | 68.8 | 84.9 | 90.8 | 99.6 |
| 3rd Quantile | 188.4 | 271.2 | 260.6 | 236.8 |
| Missing | 1 (0.2%) | 4 (0.8%) | 6 (1%) | 12 (2.6%) |
| **Hemoglobin, g/dL\*** |  |  |  |  |
| 1<sup>st</sup> Quantile | 11.6 | 11.5 | 11.7 | 11.3 |
| Median | 12.6 | 12.7 | 12.7 | 12.5 |
| 3rd Quantile | 13.6 | 13.7 | 13.5 | 13.5 |
| Missing | 3 (0.6%) | 10 (1.9%) | 0 (0%) | 4 (0.9%) |
| **Albumin, g/L\*** |  |  |  |  |
| 1<sup>st</sup> Quantile | NA | 41 | 38 | 40 |
| Median | NA | 43 | 42 | 43 |
| 3rd Quantile | NA | 45 | 45 | 46 |
| Missing | 476 (100%) | 1 (0.2%) | 16 (2.7%) | 2 (0.4%) |
| **Alkaline phosphatase, U/L\*** |  |  |  |  |
| 1<sup>st</sup> Quantile | 80 | 81 | 85 | 98 |
| Median | 113 | 124 | 135 | 155 |
| 3rd Quantile | 213 | 265 | 270 | 328 |
| **Aspartate aminotransferase, U/L** |  |  |  |  |
| 1<sup>st</sup> Quantile | 20 | 19 | 20 | 20 |
| Median | 24 | 24 | 25 | 25 |
| 3rd Quantile | 31 | 31 | 33 | 33 |
| Missing | 4 (0.8%) | 1 (0.2%) | 8 (1.3%) | 3 (0.6%) |
| Data are quantiles (1<sup>st</sup>, median, 3<sup>rd</sup>) or n (%). ECOG PS=ECOG Performance Status, LD=Lactate dehydrogenase, PSA= Prostate-Specific Antigen. Albumin for ASCENT2 was missing and LDH tests for VENICE were almost all missing. * represent variables that are significantly different between the trials. |  |  |  |  |
| **Table 1. Baseline characteristics** |  |  |  |  |

**Figure 1.**
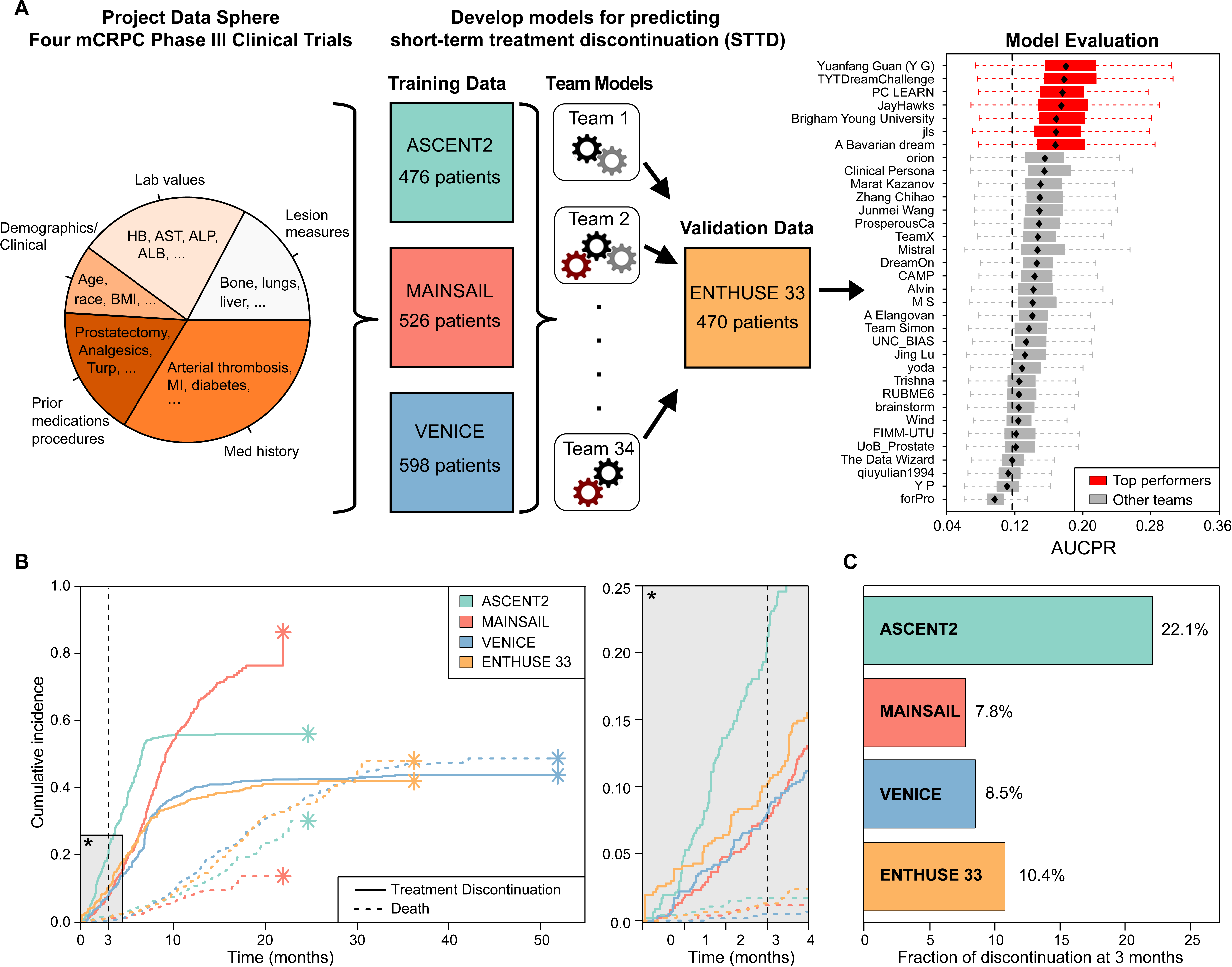
Study design and treatment discontinuation across trials. (A) Data was acquired from PDS and centrally curated by the organizing team to create a standardized dataset across the four studies. Three of the studies (ASCENT2, VENICE, MAINSAIL) were selected as training sets, and a fourth dataset (ENTHUSE 33) was withheld as a validation set. Teams submitted risk scores for evaluation in the validation set, which were scored and ranked using the area under the precision recall curve (AUPRC). (B) Trial-specific cumulative incidence functions for treatment discontinuation due to adverse or possible adverse events (solid lines) and death (dotted lines). (C) Fraction of mCPRC cases that discontinued treatment less than or equal to three months after initiation due to adverse or possible adverse events.

In total, 61 submissions were received from 34 independent, international teams participating in this Challenge. A summary of each team’s approach to data processing, handling of missing data, and statistical modeling is given in Table S3. Among teams responding to a post-Challenge survey, the five most common clinical features used in prediction models were hemoglobin (HB), alkaline phosphatase (ALP), aspartase aminotransferase (AST), prostate specific antigen (PSA), and ECOG performance status (Figure 2).

**Figure 2.**
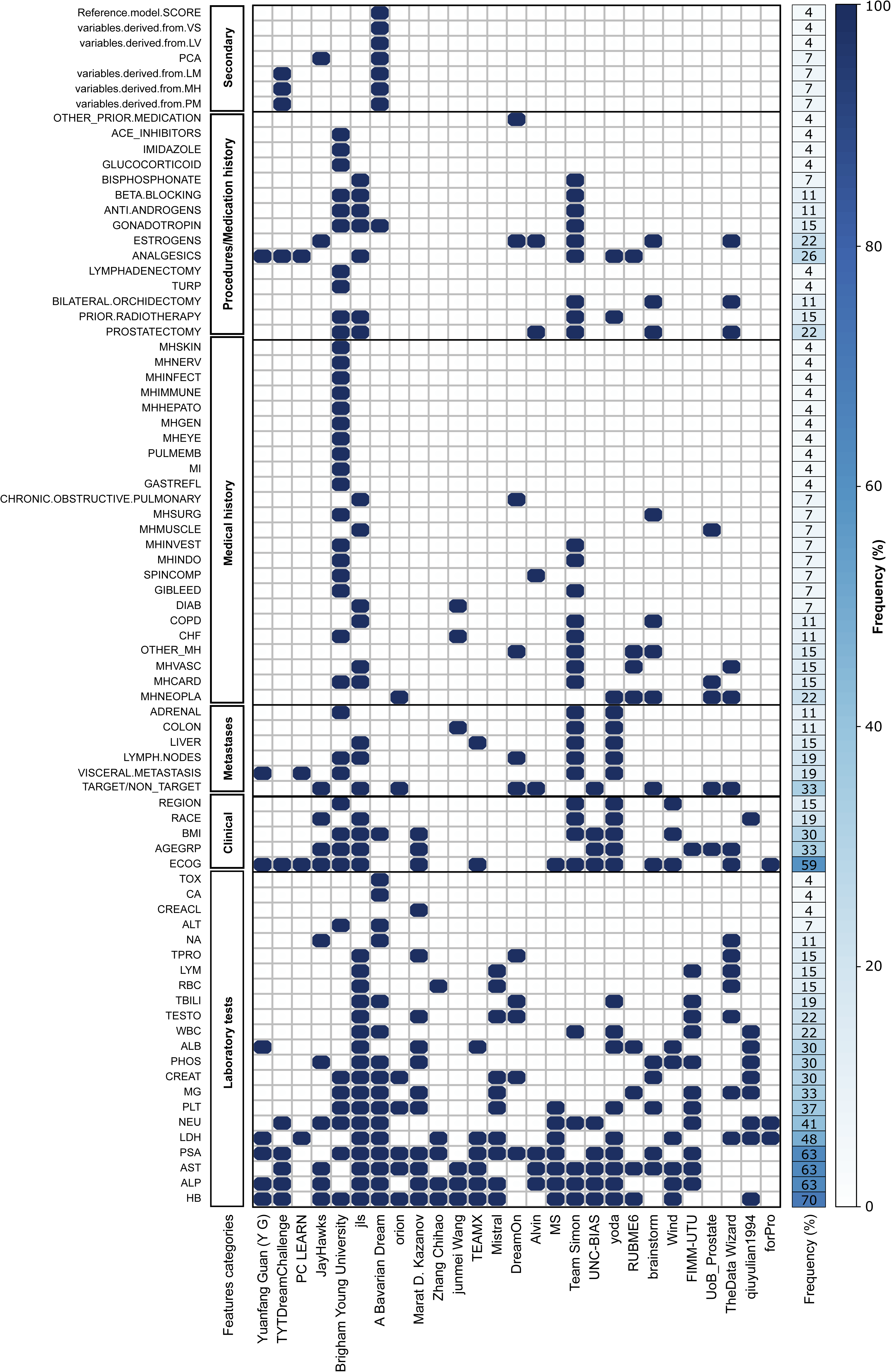
Most frequent clinical features used in prediction models. The abbreviated terms are given in Table S6.

The scoring metric of AUPRC was selected to focus on the prediction of patients that discontinued treatment, which is roughly 10% of the overall population. Across all submissions, AUPRC ranged between 0.088 and 0.178, with 0.104 representing the expected AUPRC for a random prediction model, reflective of the ~10% rate of discontinuation (Figure 1A, Table S4, Fig. S4). Team *Yuanfang Guan* (*Y G*) recorded the top score, however six other teams: *TYTDreamChallenge, PC LEARN, JayHawks, Brigham Young University, jls*, and *A Bavarian Dream*, achieved AUPRCs that were within a Bayes factor of three when compared to team *Yuanfang Guan* (Table S4, Fig. S4). While 30 out of 34 teams submitted models with potential clinical value, achieving a better AUPRC than what would be expected at random, only the previously named seven teams achieved AUPRCs that represented a statistically significant improvement over a random prediction model (adjusted *P* < 0.10) (Table S4). Consequently, these seven teams were identified as the Challenge top performers.

A cumulative lift chart analysis was performed on each submission to demonstrate the clinical utility of prediction models and to provide a more meaningful context for their associated risk predictions. Across models, area under the lift ratio curves ranged from 0.77 – 1.40 (Table S5) with an average value of 1.17; that is, prediction models improved the identification of early discontinuation events by 17%, on average, when compared to a situation where no such model(s) are used to inform patient risk. By comparison, the average area under the lift ratio curve was 1.34 among the seven top performers, representing a two-fold increase in the ability to accurately identify short-term discontinuation events compared to the average across all Challenge submissions (34% versus 17%). Restricting the above analysis to patients with high-predicted risk (i.e., top 10% of patients with highest predicted risk) revealed that models submitted by seven top performers improved the identification of early discontinuation events by a factor of two, on average, when compared to a situation where no such model(s) were used to inform patient risk (Table S5).

To understand similarities and differences in the risk predictions generated by the top performers, we hierarchically clustered patients in the ENTHUSE 33 trial data sets using the ranked patient risk scores computed from the seven top performing teams’ models. This analysis resulted in three clusters/groups of patients: patients that were consistently predicted to have a high-risk of early discontinuation (concordant high-risk; *n* = 50), patients consistently predicted to have a low-risk of early discontinuation (concordant low-risk; *n* = 170), and a group of patients with discordant risk scores across the top performers (discordant risk; *n* = 234 patients) (Figure 3A). Notable variation in the cumulative incidence of short-term treatment discontinuation events was observed between the three groups, with the concordant high-risk group exhibiting a nearly two-fold increased proportion of discontinuation events at three months compared to the concordant low risk and discordant groups (Figure 3B). Specifically, at three months post-treatment, 26% of the patients in the concordant high-risk cluster discontinued docetaxel, compared to only 9% in both the concordant low-risk and discordant groups. In addition, the competing-risk (i.e. death) was considerably elevated in the concordant high-risk cluster (6% death rate at three months) compared to the concordant low-risk and discordant groups; zero deaths observed at three months in the latter two groups.

**Figure 3.**
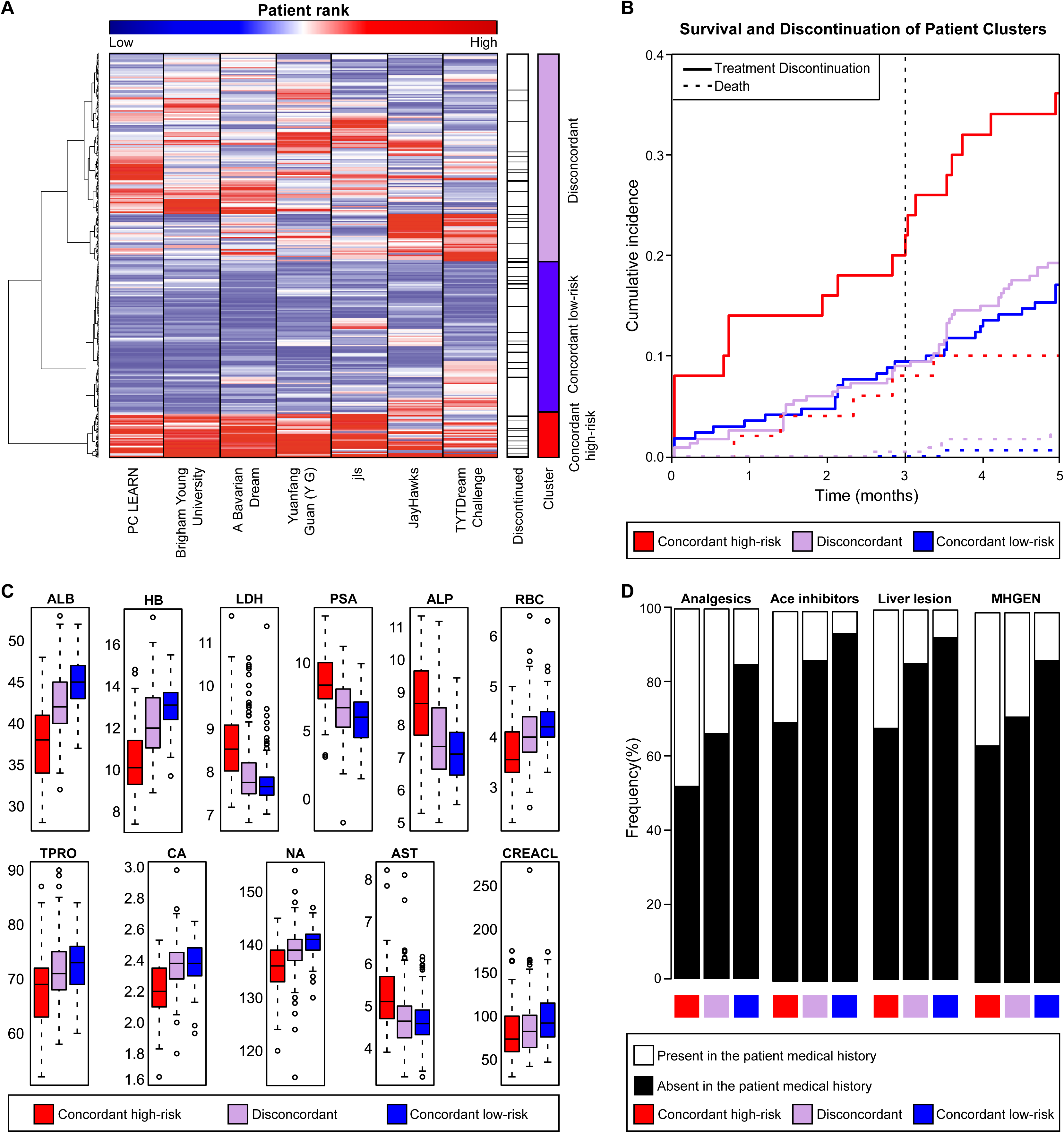
Meta-analysis of risk scores computed by the seven top performing teams. (A) Hierarchical clustering heat map of patients in the ENTHUSE 33 validation data set (*n* = 470) based on their normalized ranked risk score, computed across the seven top performing teams. (B) Kaplan Meier curves, stratified by event type (i.e., death or treatment discontinuation) across the three identified patient subgroups. (C) Distribution of baseline lab variables found to be significantly different between the three patient subgroups. (D) Distribution of baseline prior medical and medication variables found to be significantly different between the three patient subgroups. Abbreviated terms are given in Table S6.

A comparison of baseline characteristics across the three groups revealed eleven statistically significant lab values (adjusted *P* < 0.05), including: albumin (ALB), hemoglobin (HB), lactate dehydrogenase (LDH), prostate specific antigen (PSA), sodium (NA), red blood cell (RBC), alkaline phosphatase (ALP), calcium (CA), aspartase aminotransferase (AST), creatinine clearance (CREACL), and total protein (TPRO) (Figure 3C). In addition, ECOG performance status, metastatic liver lesions, and use of analgesics and ace inhibitors differed significantly between the concordant high- and low-risk clusters (adjusted *P* < 0.05) and use of analgesics and ACE inhibitors was significantly elevated among patients in the concordant high-risk cluster (48% and 30%, respectively) compared to those in the concordant low-risk cluster (15% and 5%, respectively) (Figure 3D). A similar trend was observed in the frequency of patients with liver metastasis; liver lesions were reported for only 8% of patients in the concordant low-risk cluster compared to 32% in the high-risk cluster.

Motivated by the “wisdom of crowds” performance seen in previous Challenges and the modest correlation of the risk scores across the seven top performers (Fig. S5), we aimed to determine if further improvements to prediction accuracy were possible by combining individual models submitted to the Challenge. After completion of the Challenge, we developed a community-based, ensemble classifier as a weighted function of the risk scores generated from the top seven performing teams’ models. Weights were empirically determined, and proportional to a model’s performance when evaluated in a randomly selected subset of the Challenge training data set (Fig. S2). The ENTHUSE 33 data remained an entirely independent data set for benchmarking the prediction performance of Challenge submissions, including the ensemble model. Application of the ensemble-based model to the ENTHUSE 33 resulted in an AUPRC of 0.230, outperforming the top Challenge submission by a margin of 0.052, which exceeds the difference in AUPRC between the Challenge top performers and the next best Challenge submission (Figure 1A and Figure 4A). In repeated bootstrap sampling of ENTHUSE 33 data set, the ensemble-based model outperformed the Challenge top performers the majority of times (73.4% to 94.7% across the top seven models), and achieved a Bayes factor > 3 when compared to all but a single Challenge submission; team *Yuanfang Guan* being the exception, with a Bayes factor of 2.75 (Fig. S6). The Bayes factor results reflect a direct comparison between two methods evaluated using random samplings of the ENTHUSE 33 dataset, where a Bayes factor of 3, for example, means that the first method outperformed the second method at a ratio of 3:1.

**Figure 4.**
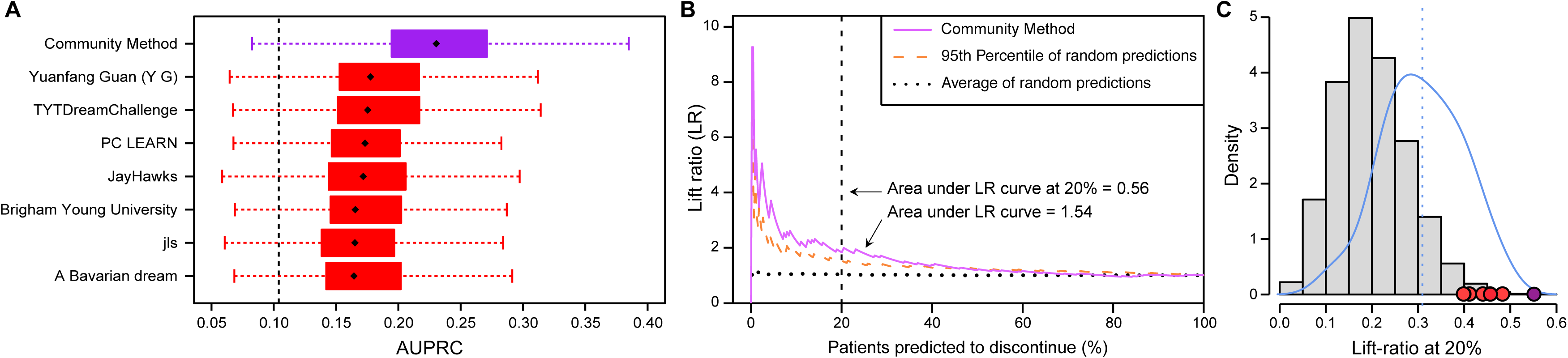
Performance of the post-Challenge ensemble-based prediction model. (A) Area under the precision recall curve (AUPRC) computed within the ENTHUSE 33 data set for the ensemble-based prediction model, along with the models developed by the seven top performing teams. Black diamonds represent the observed AUPRCs and horizontal boxplots reflect the empirical distribution of a model’s AUPRC based on 5,000 bootstrap samples generated from each models’ predictions. Vertical dotted line represents the mean AUPRC computed from 5,000 bootstrap samples generated from a random prediction model. (B) Lift-ratio (LR) curve for the ensemble-based prediction model with grey lines representing the LR-curves generated for 100 random prediction models. (C) Distribution of the area under the LR curve at 20% based on random prediction models (grey), all challenge submissions teams (blue), the top-performing teams (red and orange points), and the post-challenge ensemble-based classifier (purple).

A cumulative lift chart analysis of risk predictions computed from the ensemble-based model showed a 14% improvement (in absolute percentage points) over the top Challenge submission for correctly identifying patients that discontinued docetaxel treatment within three months (Figure 4B). Further analysis revealed a statistically significant increase in the area under the lift ratio curve at 10% generated using risk predictions from the ensemble-based method (*P* < 0.01) (Figure 4C).

To understand our ensemble-based prediction model within the broader context of clinical trial design, we conducted a simulation study to compare the sample size requirements of clinical trials that incorporate risk estimates for early treatment discontinuation to inform patient inclusion within the treatment arm. The results of our simulation study showed that when patient selection into the trial is completely random (invariant with respect to risk for early treatment discontinuation), the sample size required for detecting a HR = 1.30 between treatment arms at 80% statistical power and at type 1 error rate of 5%, was n= 1,548 (averaged across 100 simulated data sets)(Fig. S7). However, when selection into the trail is based on a patient’s risk for early treatment discontinuation, the estimated sample size required for detecting a HR = 1.30 when the accuracy for correctly identifying patients that discontinue treatment early was consistent with the performance of the ensemble-based model, was *n* = 1,306 (averaged across 100 simulated data sets)(Fig. S7); a reduction of 242 fewer patients. Simulation results across a range of prediction accuracies can be found in Fig. S7.

Acknowledging that the clinical utility of the ensemble-based prediction model is limited by the fact that several of its constituent models (i.e., models developed by the seven top-performers) used many, and in some cases all of the baseline clinical features to inform risk predictions, we developed a second, parsimonious prediction model using a restricted subset of baseline clinical features that discriminated high-versus low-risk patients for early treatment discontinuation to arrive at 5 variables: HB, ALB, PSA, NA, and LDH(Fig. S8). Prediction performance in the validation data set was comparable to the performance achieved by the ensemble-based prediction model (AUCPR = 0.236) despite using many fewer baseline clinical features. We have created a publicly available web-based implementation of this model, which can be freely accessed at the following weblink: <u>http://dream.web.tool.aicml.ca/</u>

## Discussion

The clinical value of prediction models for early treatment discontinuation on the basis of a patient’s clinical characteristics is now widely recognized and supported by a growing number studies that have begun to address this problem for a range of different disease-treatment combinations^27–29^. In the absence of effective models for predicting early failure of docetaxel treatment, many clinicians will instead use factors associated with poor survival outcomes to guide treatment decisions; for example, identifying candidates for a docetaxel treatment regimen based on an assessment of a patient’s long-term prognosis. These risk factors typically include: ALP, HB, ALB, PSA, LDH, ECOG PS, disease site (divided into three categories of lymph node only, bone/bone + lymph node, or any visceral) and use of analgesics, according to a currently available model^10^. Using the results from the top seven teams, we confirmed that these variables are predictive of poor prognosis and we also discovered several other clinical variables, including PSA, RBC, CA, AST, CREACL, and TPRO, which were significant predictors of membership in the concordant high-versus low-risk groups. Hematologic parameters and patient performance status have been previously reported as significant predictors of severe AE in patients with advanced stage non-small cell lung cancer treated with first-line chemotherapeutics^39^. Interestingly, aspartate aminotransferase (AST) was used in many of the top prediction models and was found to be significantly elevated in the high-versus low-risk groups. While further investigation is needed to understand the clinical and biological implications of these relationships, our results underscore the interrelated nature of risk predictors and the difficulty associated with finding features that are specific for toxicity-induced treatment failure.

Although seven out of 34 participating teams (20.5%) submitted models that performed significantly better than what would be expected at random when evaluated in an independent validation set, the prediction performance – even among top models – showed only moderate accuracy for correctly identifying high-risk patients for early discontinuation; on average, the top performing models resulted in a modest 34% improvement in the ability to correctly identify short-term discontinuation events compared to a situation where no such models are used. The Challenge results served to initiate post-Challenge community collaborations between Challenge organizers and members from each of the top performing teams, which aimed to improve prediction performance by leveraging the wisdom of crowds. This community effort led to the development of an ensemble-based prediction model (generated using the top performing teams’ models) that recorded the best overall prediction performance in the ENTHUSE 33 data set, and in all but a single instance, significantly better performance over individual models submitted to the Challenge. To make our finding easily accessible, we leveraged the community insight to select variables to build a webtool that can predict the risk of patient discontinuation based on 5 clinical variables: http://dream.web.tool.aicml.ca/. Our findings reinforce the idea that the collective wisdom of crowds can be effectively harnessed to produce model(s) whose predictive value exceeds that obtained by individual members of the crowd^40, 41^. Further, these results establish a precedent for combining models in future crowdsourced challenges.

Although the post-Challenge ensemble-based prediction model lacks the accuracy needed for immediate clinical application^42^, this study is nevertheless a critical first step in the development of viable clinical tools and is the first to establish a performance benchmark for future prediction models of this sort. Importantly, our findings have the potential to immediately impact future mCRPC clinical trials with a docetaxel-based treatment arm by improving patient selection through the use of novel selection designs. Indeed, we showed through a simulation study that effective prediction of patients that will discontinue due to adverse events can reduce patient enrollment by significant numbers, especially when the difference between controls and treatment is low. While future work would be needed to investigate how to best integrate the models described here in the context of these and/or other designs, the prospect is encouraging and inline with a growing emphasis on the need for innovative approaches for clinical trial design^44^.

Notwithstanding its highlights, there are several limitations associated with this work. Since the initiation of the four trials used in this Challenge, several promising therapies have emerged that have reshaped the treatment of mCRPC^45^. While changing treatment paradigms may limit the generalizability of the prediction models reported here, the fact that several predictors of early docetaxel discontinuation coincided with previously identified markers of poor-prognosis point to the existence of a general class of prognostic/predictive features in the context of mCRPC patient outcomes. This class of clinical features may therefore serve as a useful starting point for future studies focused on the identification early discontinuation risk predictors for new and emerging treatment regimens. A second limitation of this study is that there was no restriction imposed by Challenge organizers on the number of clinical features used in the development of prediction models. As a result, several of the prediction models submitted to the Challenge (including the post-Challenge ensemble-based model) used many or all of the baseline clinical features contained in the standardized data table. While this may create challenges for future studies seeking an internal comparison of model performance metrics (i.e., side-by-side comparison of AUPRC) in data sets other than those used in this study, the AUPRCs reported here can nevertheless be used as benchmarks to gauge the performance models developed and evaluated in other data sets.

The DREAM Challenge described here exemplifies how open-access cancer trial data can be used to explore new clinical questions and highlights the role of crowdsourcing as a tool for advancing predictive models for cancer outcomes. The Challenge has also demonstrated the willingness of the research community to work together to advance predictive modeling in mCRPC. Strikingly, the group of researchers that performed the post-Challenge analysis, developed the ensemble predictor, and wrote this manuscript had never worked together before. The challenges we face in biomedical science are too great for siloed research to be the status quo moving forward. Fostering research in this manner is further evidence that the biomedical research of tomorrow can and will be a team effort.

## Acknowledgements

This publication is based on research using information obtained from <u>http://www.projectdatasphere.org/</u>, which is maintained by *Project Data Sphere, LLC* (PDC). Neither PDC, nor the owner(s) of any information from the website, have contributed to, approved, or are in any way responsible for the contents of this publication. We thank the Sage Bionetworks Synapse team for the development and design of the Challenge website. This work is supported in part by the following: National Institutes of Health, National Library of Medicine (2T15-LM009451), National Cancer Institute (5R01CA152301), Boettcher Foundation, Doctoral Programme in Mathematics and Computer Sciences at the University of Turku, European Union’s Horizon 2020 research and innovation programme, Academy of Finland, Juvenile Diabetes Research Foundation JDRF, and Sigrid Juselius Foundation.

## Declaration of interests

Dr. Sweeney reports personal fees from Sanofi, personal fees from Janssen, personal fees from Astellas, personal fees from Bayer, outside the submitted work; Dr. Zhou reports employment and stocks from Sanofi US, outside the submitted work; Dr. Shen reports employment and stocks from Sanofi US, outside the submitted work; Dr. Abdallah reports employment and stocks from AstraZeneca, outside the submitted work; Dr. Scher reports non-financial support from Astra Zeneca, personal fees from Astellas, personal fees from BIND Pharmaceuticals, personal fees from Blue Earth Diagnostics, non-financial support from Bristol Myers Squibb, personal fees from Clovis Oncology, personal fees from Elseiver’s PracticeUpdate Website, non-financial support from Ferring Pharmaceuticals, personal fees from Genentech, personal fees from Med IQ, non-financial support from Medivation, personal fees from Merck, personal fees from Roche, personal fees from Sanofi Aventis, non-financial support from Takeda Millennium, personal fees from WCG Oncology, personal fees from Asterias Biotherapeutics, grants from Illumia, Inc, grants from Innocrin Pharma, grants from Janssen, grants from Medivation, outside the submitted work; MSc. Seyednasrollah reports grants from Doctoral Programme in Mathematics and Computer Science at the University of Turku, grants from Sigrid Jeselius Foundation, during the conduct of the study; Dr. Sartor reports grants and personal fee from Sanofi, outside the submitted work; Dr. Elo reports grants from European Research Council (ERC), European Union’s Horizon 2020 research and innovation programme, Academy of Finland, Juvenile Diabetes Research Foundation JDRF, and Sigrid Juselius Foundation, during the conduct of the study; The other authors declared no conflicts of interest.

## Author Contributions

T.W., C.B., E.C.N., T.Y., K.A., T.N., G.S., H.S., C.J.S., C.J.R., H.I.S., O.S., F.L.Z., J.G., and J.C.C. designed the Challenge. F.L.Z. and L.S. led the PDS efforts to collect and process the clinical trial data. F.S., D.C.K., T.W., S.R.P., R.V., R.G., C.F., E.G., L.K., R.D.W., K.K.W., L.L.E., F.L.Z, J.G., and J.C.C. performed the post-Challenge data analysis and interpretation. H.S., C.J.S., C.J.R., H.I.S., and O.S. assisted in clinical variable interpretation and manuscript preparation. All members of the Prostate Cancer Challenge DREAM Consortium submitted prediction models to the Challenge, provided method write-ups, and the code to reproduce their predictions. F.S., D.C.K., T.W., S.R.P., R.G., C.F., R.D.W., H.S., C.J.S., C.J.R., H.I.S., O.S., L.L.E., F.L.Z., J.G., and J.C.C. wrote the manuscript.

